# Gene conversion explains elevated diversity in the immunity modulating APL1 gene of the malaria vector *Anopheles funestus*

**DOI:** 10.1101/2021.11.25.470000

**Authors:** Jack Hearn, Jacob M. Riveron, Helen Irving, Gareth D. Weedall, Charles S. Wondji

## Abstract

The leucine rich repeat gene APL1 is a key component of immunity to *Plasmodium* and other microbial pathogens in *Anopheles* mosquitoes. In the malaria vector *Anopheles funestus* the APL1 gene has four paralogues which occur along the same chromosome arm. We show that APL1 has exceptional levels of non-synonymous polymorphism across the range of *An. funestus* with an average π_n_ of 0.027 versus a genome-wide average of 0.002, and π_n_ (and π_s_) is consistently high in populations across Africa. The pattern of APL1 diversity was consistent between independent pooled-template and target-enrichment datasets, however no link between APL1 diversity and insecticide-resistance was observed with the phenotyped target-enrichment dataset. Two further innate immunity genes of the gambicin anti-microbial peptide family had π_n_/π_s_ ratios greater than one, possibly driven by either positive or balancing selection. Cecropin antimicrobial peptides were expressed much more highly than other anti-microbial peptide genes, an observation discordant with current models of anti-microbial peptide activity. The observed APL1 diversity likely results from gene conversion between paralogs, as evidenced by shared polymorphisms, overlapping read mappings, and recombination events among paralogues. Gene conversion at APL1 versus alternative explanations is concordant with similarly elevated diversity in APL1 and TEP1 loci in *An. gambiae*. In contrast, the more closely related *An. stephensi* which also encodes a single-copy of APL1 does not show this elevated diversity. We hypothesise that a more open chromatin formation at the APL1 locus due to higher gene expression than its paralogues enhances gene conversion, and therefore increased polymorphism, at APL1.

## Introduction

In 2018, there were 228 million cases of malaria worldwide, leading to 405,000 deaths. The majority of cases (93%) and deaths (94%) occurred in Sub-Saharan Africa, and *Plasmodium falciparum* was overwhelmingly responsible (>99%) (WHO 2020). *P. falciparum* is vectored by *Anopheles* mosquitoes, and vector competence (susceptibility to *Plasmodium*) varies greatly between species due to the action of immune genes (White et al. 2011; Blandin et al. 2009). Furthermore, rising insecticide resistance may increase or decrease vector competence of *Anopheles* mosquitoes by stimulating or blocking mosquito immune responses (Rivero et al. 2010).

Insects lack the adaptive immune response of vertebrates, relying entirely on a powerful innate immune system to fight pathogens. Mosquitoes are no exception, employing a complement-like system acting in the haemolymph that identifies and targets pathogens for destruction (Clayton et al. 2014). In *Anopheles* species the complement-based response to *Plasmodium* and pathogenic microbes is mediated by a dimer of *Anopheles Plasmodium*-responsive Leucine rich repeat protein 1 (APL1) and Leucine Rich Immune Molecule 1 (LRIM1), which form a complex with thioester-containing protein 1 (TEP1) (Baxter et al. 2007) a homologue of the vertebrate complement protein C3. Together, this complex recognises pathogens of bacterial and fungal origin in addition to the ookinete stage of *Plasmodium*, and triggers a cascade that eliminates them through lysis or melanisation (Povelones et al. 2013; Reyes Ruiz et al. 2019). Another component of Anopheles innate immune systems are anti-microbial peptides (AMPs) which have a range of antibacterial, antifungal and antiviral activities (Wu et al. 2018). AMPs are split into five families, which are variable in presence and copy number across mosquito species (Lee et al. 2019). A cysteine-rich AMP named gambicin was marginally lethal to *P. berghei* ookinetes (Vizioli et al. 2001). Despite these defence mechanisms, *P. falciparum* is still well able to evade detection by mosquito immunity complexes through the action of *Pfs47* surface protein and mosquito receptor in a ‘lock and key’ fashion (Molina-Cruz et al. 2013; Cha & Jacobs-Lorena 2020).

Among *Anopheles* species, *An. gambiae* and *An. coluzzii* have the best studied set of APL, LRIM and TEP genes, and silencing any one of these genes in these species enhances *Plasmodium* infectivity (Molina-Cruz et al. 2012; Povelones et al. 2009). Of particular note is the presence of *Plasmodium*-resistant strains of *An. gambiae s. l.* due to the action of these genes (Niaré et al. 2002; Riehle et al. 2006), alongside elevated genetic diversity and selective sweeps in APL1 and TEP1 genes (Rottschaefer et al. 2011; Obbard et al. 2008). The APL1 locus has duplicated to form three copies (labelled A, B, and C) in the lineage ancestral to the *An. gambiae* complex and *An. christyi* (Mitri, Bischoff, Eiglmeier, et al. 2020), but remains a single locus in other *Anopheles* species including *An. funestus*. In *An. stephensi* RNAi-mediated silencing of the single-copy APL1 results in increased mortality which is rescued by antibiotics (Mitri, Bischoff, Eiglmeier, et al. 2020). Specifically, APL1 in *An. stephensi* modulates abundance of *Klebsiella* and *Cedecea* bacteria of the Enterobacteriaceae family (Mitri, Bischoff, Belda Cuesta, et al. 2020), both of which are naturally found in *Anopheles* microbiomes. Reyes Ruiz et al (2019) proposed two models for how the APL1/LRIM1/TEP1 complex acts to eliminate pathogens. In the first model, the complex acts as a recognition molecule that changes conformation on pathogen-binding which activates a zymogen protease; in the second model, the pathogen surface is recognised by another molecule that activates a protease of haemolymph APL1, leading to the release of a processed form of TEP1 in the pathogen’s vicinity (Reyes Ruiz et al. 2019).

Exceptional polymorphism and maintenance of alleles at the APL1 and TEP1 loci in the *An. gambiae* complex were consistent with gene conversion versus other explanations like balancing selection (Rottschaefer et al. 2011; Obbard et al. 2008). Although balancing selection is thought to explain similarly raised levels of genetic diversity seen in AMPs of four populations of *Drosophila melanogaster* (Chapman et al. 2019) and may be acting on gambicin in *An. gambiae* (Lehmann et al. 2009). The molecular diversity and expression dynamics of AMP genes in *An. funestus* has not been explored to date. By contrast to APL1 and TEP1, the third component of the complement complex, LRIM1, has not been associated with higher diversity in *An. gambiae* (Obbard et al. 2008; Slotman et al. 2007). An adaptive role for APL1 and TEP1 elevated diversity may reflect protection against a variety of microbial pathogens that vary across species’ ranges and seasonality (Rottschaefer et al. 2011; Mitri, Bischoff, Eiglmeier, et al. 2020). High diversity of APL1 is not a genus-wide trait however, in *An. stephensi* the single-copy of APL1 exhibits low genetic diversity in an Iranian-sampled population (Mitri, Bischoff, Eiglmeier, et al. 2020). This could reflect a locally stable microbiome of enteric taxa that places APL1 under purifying selection. In *An. stephensi* APL1 has a dual function against *Cedecea* and *Klebsiella* Enterobacteria, and *Plasmodium,* which was demonstrated with *P. yoelii* following depletion of APL1 (Mitri, Bischoff, Eiglmeier, et al. 2020; Mitri, Bischoff, Belda Cuesta, et al. 2020). *An. gambiae sensu lato* differs as the three APL1 copies protect against *Plasmodium* infection, but not pathogenic bacteria (Mitri, Bischoff, Eiglmeier, et al. 2020) meaning an alternative mechanism must perform anti-bacterial responses.

*Anopheles funestus* is one of the major vector species of *P. falciparum* in sub-Saharan Africa. It is widespread across Sub-Saharan Africa, and in several regions it is the dominant vector of malaria (Gillies et al. 1968; Coetzee & Fontenille 2004). As a member of the *An. funestus* group it is phylogenetically distant from *An. gambiae*, *An. arabiensis*, and *An. coluzzii* of the *An. gambiae s. l.* species complex. Historically, it has been more difficult to breed and maintain laboratory colonies, until the introduction of forced-egg laying techniques (Morgan et al. 2010). Together, these factors have left *An. funestus* less well-studied in many aspects of its biology relative to the other malaria vector species. Indeed, a chromosome-level genome assembly became available much later than *An. gambiae s. l.* (Ghurye et al. 2019). Despite this, the molecular basis of resistance to pyrethroid insecticides is, arguably, best understood in this species with several metabolic resistance-conferring variants identified and functionally validated in cytochrome p450 and glutathione-S-transferase (GST) genes using both whole genome sequencing and targeted enrichment sequencing approaches (Weedall et al. 2019; Riveron et al. 2014; Mugenzi et al. 2019). Over-expression of GSTs creates a possible point of cross-talk between resistance and immunity, as over-expressed GSTs may neutralise reactive oxidative species (ROS), which are a key component of insect immune response to *Plasmodium* (Tchouakui et al. 2019; Rivero et al. 2010). There may also be an energetic trade-off between overexpression of metabolic-resistance genes like GSTs and robustness of immune responses to *Plasmodium* and other pathogens (Tchouakui et al. 2019; Rivero et al. 2010). *An. funestus* encodes single-copy orthologues of the APL1, LRIM1, and TEP1 complement genes with paralogues for each gene (VectorBase genome annotation version 51). Particularly so for the TEP-family genes, which may reflect a Diptera-wide trend for expansion in this family (Viljakainen 2015; Sackton et al. 2017). The *An. funestus* genome also encodes 11 AMP genes of the defensin, cecropin, attacin and gambicin families, but lacks diptericin gene (in annotation version 51) (Ghurye et al. 2019).

Here, we assessed the genetic diversity of the APL1/TEP1/LRIM1 and AMP innate immunity genes in *An. funestus* using a combination of genomic and expression datasets. Among AMP genes we find that two gambicin genes have elevated π_n_ /π_s_ ratios concordant with ongoing selection alongside high expression of cecropins across *An. funestus* populations. Principally, we show that the key *Plasmodium* and gram-negative bacteria response gene APL1 has high non-synonymous (π_n_) and synonymous (π_s_) diversities versus genome-wide averages for this species. Multiple sources of evidence including read mappings, shared polymorphisms, and recombination between APL paralogues suggest non-homologous gene conversion plays a key role in elevated diversities.

## Methods

### Identifying APL, LRIM, TEP and AMP genes in *An. funestus*

We selected immune genes that were directly annotated as APL, LRIM, TEP and AMP genes (a) from the *An. funestus* AfunF3.1 gene-set, (b) genes that were designated as orthologues of *An. gambiae* (AgamP1.10) genes of each category in VectorBase and (c) *An. funestus* paralogues for genes selected by criteria (a) and (b). VectorBase defines orthology and paralogy using OrthoMCL (Li et al. 2003)

### Variant prediction from genome-wide pooled-sequencing

Read data for pooled-template whole genome sequencing (POOLseq), target-enriched individual genome sequencing (SureSelect) and RNA-sequencing (RNAseq) analyses is available in the European Nucleotide Archive under accessions PRJEB13485, PRJEB24384, PRJEB35040, PRJEB24351, PRJEB24520, PRJEB47287, PRJEB48958 and PRJEB24506 (Weedall et al. 2019; Mugenzi et al. 2019; Weedall et al. 2020; Hearn et al. 2021). Pooled-sequencing data for ten F_0_ populations of *An. funestus* from eight locations and two lab strains (Weedall et al. 2019, 2020) was aligned to the AfunF3.1 genome assembly (Ghurye et al. 2019) with BWA (0.7.17) (sampling locations and alignment metrics, Supplementary Table 1). Alignment bam files were sorted, duplicates removed with Picard (2.18.15-0) (2019) and converted to mpileup format in Samtools (1.9) (Li et al. 2009). Variants were identified with Varscan (mpileup2cns, version 2.4.3) (Koboldt et al. 2009, 2012) with a p-value threshold of 0.05 and a minimum allele frequency of 0.01. Variants were left-aligned, normalised, split into biallelic sites, and SNPs filtered within 20 bp of an indel in bcftools (1.9) (Li et al. 2009). SNP effects were predicted in SnpEff (v4.3) (Cingolani et al. 2012) using a custom *An. funestus* database created from AfunF3 genome. Population genetic parameters were estimated for annotated genes using SNPGenie (2019.10.31) (Nelson et al. 2015), filtered variants, and the AfunF3.1 genome annotation. Additional genes were noted if they had high diversities and could be linked to immune function. Per gene and exon average read coverage depths were calculated in Jvarkit (version d9efbd3, https://github.com/lindenb/jvarkit) “bamstats05.jar” from bam alignment files to compare per-gene coverages versus genome-wide averages.

### Targeted sequencing of candidate resistance genes and regions

To identify APL1 variant sites with allele frequencies significantly associated with resistant phenotypes, we analysed a targeted enrichment experiment based on genes with a potential role in insecticide resistance [see (Hearn et al. 2021) for further details]. For populations of *An. funestus* from Malawi and Uganda we selected ten mosquitoes that died after 60 minutes’ exposure to permethrin and ten mosquitoes still alive after 180 minutes’ exposure. Due to lower resistance levels in Cameroon, mosquitoes that died after 20 minutes exposure to permethrin or were alive after 60 minutes were collected. Ten individuals each from the susceptible FANG lab strain originating in Angola, and the FUMOZ pyrethroid resistant strain originating in Mozambique were also included (Hunt et al. 2005). In short, a selection of potentially resistance related genes including heat shock proteins, odorant binding proteins, detoxification genes and immune response genes and all known target-site resistance genes sequences from *An. funestus* (Hearn et al. 2021). Additionally, all genes in the major quantitative trait loci associated with pyrethroid resistance were included. These were a 120kb region BAC clone of the *rp1 (resistance to pyrethoid 1)* locus containing the major CYP6 P450 cluster on chromosome 2R and the 113kb region BAC clone sequence for *rp2* on chromosome 2L (Wondji et al. 2009). A total of 1,302 target sequences were included. Baits were designed using the SureSelect DNA Advanced Design Wizard in the eArray program of Agilent and library construction and sequencing performed by the Centre for Genomic Research (CGR), University of Liverpool, using the SureSelect target enrichment custom kit. Libraries were pooled in equimolar amounts and sequenced in 2×150bp paired-end fragments on an Illumina MiSeq with 20 samples per run (version 4 chemistry).

Alignment, sorting and duplicate removal of SureSelect data for 80 individuals was the same as for pooled-sequencing. Variants were called in freebayes (v1.3.2) and filtered for a phred-scaled quality-score greater than 20 with ’vcffilter’ of vcflib (v1.0.0) (Garrison & Marth 2012). Only genes with an average coverage over 500 calculated in Jvarkit from all pooled individuals were retained for population genetic analyses. Diploid SureSelect variant data was then phased using WhatsHap (Patterson et al. 2015). Phased SNP-only genotypes for 160 haploid genomes were converted into fasta sequences in bcftools. Immune genes of interest were extracted from haplotype genomes using GffRead (Pertea & Pertea 2020) and aligned in muscle (Edgar 2004) to ensure correct positioning across haplotypes. Kimura’s 2-parameter distance haplotype networks were constructed in the R package pegas, as were per gene Tajima’s D estimates and p-values to indicate population structure and directional or balancing selection at each locus (Paradis 2010).

### F_st_-based associations between resistant and susceptible mosquitoes within each country

Non-synonymous and synonymous annotated SNPs for immune genes separately and the whole of chromosome 2 were input to the R package poolfstat (v2.0.0) in variant calling format (VCF) for global F_st_ estimates from African populations (Hivert et al. 2018). An F_st_ based association study was performed APL1 variants between the ten susceptible (dead) and ten resistant (alive) mosquitoes generated by the SureSelect experiment. pFst (https://github.com/vcflib/vcflib) was run on the freebayes predicted variants (applying flag “--type GL” in pF_st_). The analysis was restricted to bi-allelic coding sequence variants, and resulting p-values were false discovery rate adjusted using qvalues (Dabney et al. 2010) package in R and a threshold 0f 0.05 applied.

### Gene expression of APL1, TEP and LRIM genes

RNASeq data for 46 replicates of *An. funestus* (first published in Weedall et al., 2019) from Ghana, Uganda, Cameroon, Malawi, FANG and FUMOZ was aligned to the genome using the subread aligner (2.0.1) and quantified using featureCounts of the Subread package (Liao et al. 2013). Raw counts were converted to transcripts per million (TPM) values (following http://ny-shao.name/2016/11/18/a-short-script-to-calculate-rpkm-and-tpm-from-featurecounts-output.html) to allow comparison between genes, and average TPM per gene calculated from all replicates combined. These RNASeq data were generated from total RNA of mosquitoes that survived exposure to pyrethroids (resistant) and DDT or that were not exposed to insecticides (Control) and resistant (FUMOZ) and susceptible (FANG) laboratory strain. A differential gene expression analysis between countries (combined by insecticide treatment) and FANG and FUMOZ laboratory strains was performed on the raw counts in DESeq2 using the iDEP server (version 94) (Ge et al. 2018). Counts were filtered to include only genes with a minimum count per million of 0.5 in at least four libraries. All 15 possible pairwise contrasts were tested, with a false discovery rate cut-off of 0.05 used to accept significance. Of the 13,144 genes that passed iDEP filtering only results for immunity genes of AMP, APL, LRIM, TEP family genes and their paralogues were investigated further.

### Identifying shared polymorphisms and recombination between APL1 paralogues

Discordantly mapped read pairs, in which each read of the pair mapped to a different paralogue were quantified for each population. These discordantly mapped pairs may result from gene conversion homogenising sequences between paralogs, or if spurious, lead to inflated diversity estimates at APL1 and paralogs. Discordant mappings were removed for each paralogue by filtering unmapped, secondary, and supplementary alignments in Samtools. Variants were re-called for these genes in Varscan, annotated with SnpEff, and gene-wide diversity metrics and coverages re-inferred with SNPGenie and bamstats05 respectively. To identify shared polymorphisms in the stringently re-mapped data a codon-aware alignment of the five paralogues was created in macse2 (v2.05), and the Varscan predicted synonymous and non-synonymous SNPs per gene converted to positions in the multiple sequence alignment. VCF files for each position were then combined to identify shared polymorphisms positions. Secondly, locations of discordant mappings with predicted insert sizes greater than 10,000 bp were identified and counted for each paralogue by intersecting the read pair mapping with gene annotations in bedtools for PoolSeq and SureSelect data.

Recombination between paralogues was tested with GARD (Genetic Algorithm in Recombination Detection) on the Datamonkey Adaptive Evolution Server. GARD was run with defaults and with ‘General Discrete’ site-to-site variation and 3 rate classes to check consistency between parameters. Coiled coil domains were predicted for each paralogue using DeepCoil2 (Ludwiczak et al. 2019)

## Results

### Immune gene complements of *An. funestus*

We identified one copy of APL1 (AFUN018743) syntenic with APL1C in *An. gambiae* and four paralogues, 13 LRIM genes and 36 TEP genes in the *An. funestus* genome annotation. Their designations and chromosomal locations are given in Supplementary Table 2. *An. funestus* APL1 and the four paralogues all encode a leucine rich repeat domain and two 3’ end coiled coils similar to their orthologues in *An. gambiae*. None of the *An. funestus* APL1 genes, however, share the repeated amino acid motif ‘P-A-N-G-G-L’ present in the 5’ end of *An. gambiae* APL1C. None of the four paralogues of APL1: AFUN018581, AFUN000279, AFUN000288 and AFUN000597 are syntenic with APL1A/B/C of *An. gambiae*. All five *An. funestus* genes are orthologous to APL1B of *An. stephensi* (ASTEI02571) which lacks paralogues (Mitri, Bischoff, Eiglmeier, et al. 2020), while only APL1/AFUN018743 is syntenic. The phylogenetically closest species to *An. funestus* with VectorBase genome annotations are *An. culicafacies* and *An. minimus*. Both species encode three genes orthologous only to the five APL1-like genes of *An. funestus*. TEP and LRIM genes, TEP1 (AFUN018758) and LRIM1 (AFUN005964) orthologues were defined by synteny to their *An. gambiae* equivalents in VectorBase (AGAP010815 and AGAP006348 respectively). Most APL1, LRIM and TEP-like genes are present on chromosome 3, with eight on chromosome 2 and zero on the X chromosome.

Two anti-microbial peptide family gambicin genes (AFUN006610 and AFUN006611) are encoded in the *An. funestus* genome which occur consecutively on chromosome 2 (around position 65.13 mb). A third consecutive gene, AFUN006612, is a VectorBase paralogue of AFUN006611 but is annotated as an unspecified product. Other AMP genes identified include four cecropins, four defensins and one attacin gene, only diptericin is lacking from the *An. funestus* genome.

### APL1 has elevated diversity

The complete PoolSeq and SureSelect datasets consisted of 12,968 and 952 genes with SNPgenie polymorphism data respectively. In the PoolSeq data 229 genes had a π_N_/π_s_ ratio greater than one, 36 of which were annotated with functions including one of the gambicin genes (AFUN006610), two of the defensin genes (AFUN016516 and AFUN016588), two salivary gland proteins (AFUN016070 and AFUN016250) and a cuticular protein RR-1 (AFUN000936). Of the genes included in the targeted-enrichment data, π_N_/π_s_ was greater than one for the same gambicin, but not the salivary gland and cuticular protein genes.

In both PoolSeq and SureSelect data *An. funestus* APL1 is an outlier in diversity levels compared to genes of similar length (Nonsynonymous sites polymorphism π_N_, Figure 1) with a π_N_ of 0.027 and π_s_ of 0.036 in PoolSeq data (Table 1 and Supplementary Table 3). Both elevated against genome-wide averages of 0.002 and 0.021 for π_N_ and π_s_ respectively (Supplementary Table 4). This was consistent across populations including the insecticide susceptible lab strain FANG, with π_N_ ranging from 0.023-0.030 (Supplementary Table 4). The APL1 paralogues also had elevated diversities, although to a lesser extent (Table 1). When ranked by PoolSeq π_N_ values, APL1 had the 15 highest of the 12,968 genes included, and only three of those with greater π_N_ were of more than 400 nonsynonymous sites across the gene (Supplementary Table 3). Immunity genes of the LRIM and TEP families do not show similarly elevated diversities with highest per gene π_N_ value of 0.013 (AFUN005964/LRIM1) and 0.017 respectively (Supplementary Table 3), nor do any genes from these families have a π_N_/π_s_ ratio greater than one. The TEP gene with highest π_N_ (AFUN019003) had no synteny to other TEP genes in VectorBase. Average coverage of each gene from the PoolSeq data indicated that APL1 and paralogue AFUN018581 have elevated coverage versus genome-wide means across populations (Table 2), however coverage profiles of these genes were both lower (less than 2-fold genome averages) and dissimilar in shape to those of known *An. funestus* duplication events (Supplementary Figure 1, for comparison with a known duplication). Instead of a sharp well-defined region of doubled (or more coverage) as seen for duplications, regions of increased coverage are non-uniform across these genes. This suggests an alternative explanation such as gene conversion explains the observed coverage variability.

**Figure 1.**
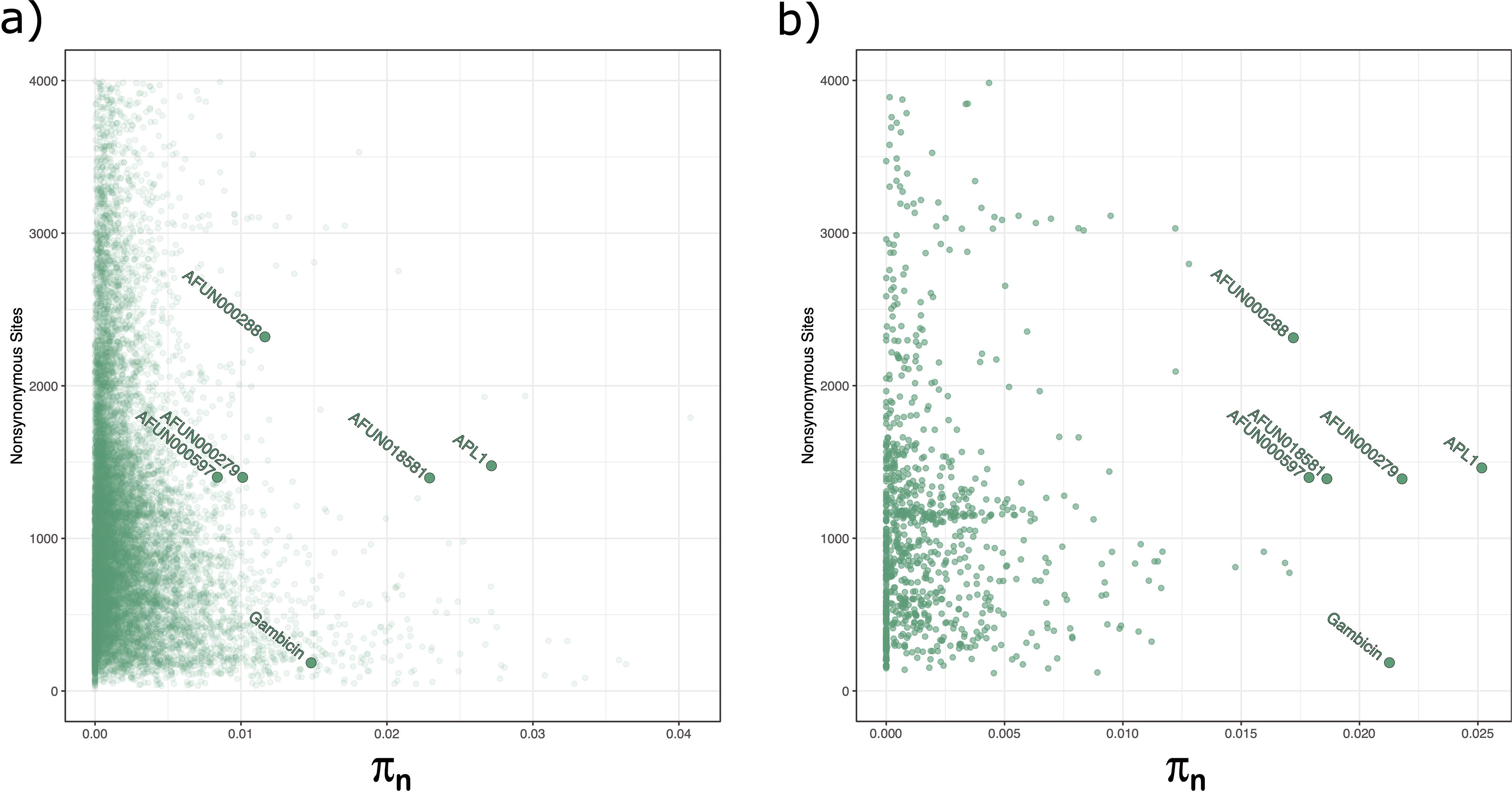
APL1 (AFUN018743) is an outlier in non-synonymous diversity. Number of non-synonymous sites versus π_N_ for all genes (a. PoolSeq data) and a selection of potentially resistance associated loci (b. SureSelect data). APL1 paralogues also have high π_N_ relative to other genes, but to a lesser extent and with more variability between data types. The anti-microbial gene gambicin (AFUN006611) is also labelled. piN = π_N._

**Table 1.** Diversity estimates for APL1 and its paralogues in *An. funestus*.

| Gene | Nonsynonymous sites | Synonymous sites | $\pi_N$ | $\pi_S$ |
| --- | --- | --- | --- | --- |
| **PoolSeq** |  |  |  |  |
| APL1/AFUN018743 | 1475.80 | 426.20 | 0.027 | 0.036 |
| AFUN018581 | 1396.35 | 409.62 | 0.023 | 0.036 |
| AFUN000279 | 1399.86 | 406.14 | 0.010 | 0.012 |
| AFUN000288 | 2320.70 | 688.30 | 0.012 | 0.019 |
| AFUN000597 | 1401.36 | 404.64 | 0.008 | 0.008 |
| **SureSelect** |  |  |  |  |
| APL1/AFUN018743 | 1461.21 | 418.62 | 0.025 | 0.027 |
| AFUN018581 | 1389.97 | 402.91 | 0.019 | 0.023 |
| AFUN000279 | 1389.10 | 402.13 | 0.022 | 0.028 |
| AFUN000288 | 2313.82 | 689.23 | 0.017 | 0.025 |
| AFUN000597 | 1398.40 | 404.38 | 0.018 | 0.020 |

Nonsynonymous sites = number of nonsynonymous positions across gene;
Synonymous sites = number of synonymous positions across gene; $\pi_N$ = nucleotide858 diversity at nonsynonymous sites; $\pi_S$ nucleotide diversity at synonymous sites.

**Table 2.** Average gene coverages from combined PoolSeq data versus background genome-wide averages for APL1 and its paralogues.

| Gene | Original Coverage | Discordant read filtered coverage |
| --- | --- | --- |
| AFUN018743 | 48.31 | 42.79 |
| AFUN018581 | 49.55 | 43.16 |
| AFUN000279 | 20.79 | 17.06 |
| AFUN000288 | 23.12 | 20.48 |
| AFUN000597 | 14.60 | 12.15 |
| All genes | 33.98 | N/A |

### Haplotype analyses and F_st_ between countries reveals no clustering by resistance or origin

Haplotype analyses of APL1 (Figure 2) revealed a high degree of intermingling among regions and resistance with no discernible groupings. Most haplotypes (145/160) were unique and did not group with any other sequences, the largest cluster was of six FANG sequences, followed by a cluster of three also FANG sequences. TEP1 was very similar in an overall lack of shared haplotypes with 144 of 160 being unique. Only two clusters of eight sequences each, both consisting entirely of FUMOZ sequences were present. For both genes mutational separation between haplotypes was high (represented by circles on connecting nodes in Figure 2). LRIM1 by contrast only presented two clusters across all 160 haplotypes, one of 152 sequences and on of 8 which were separated by only one mutation. Tajima’s D tests for detecting population structure or selection were non-significant for all three genes.

**Figure 2.**
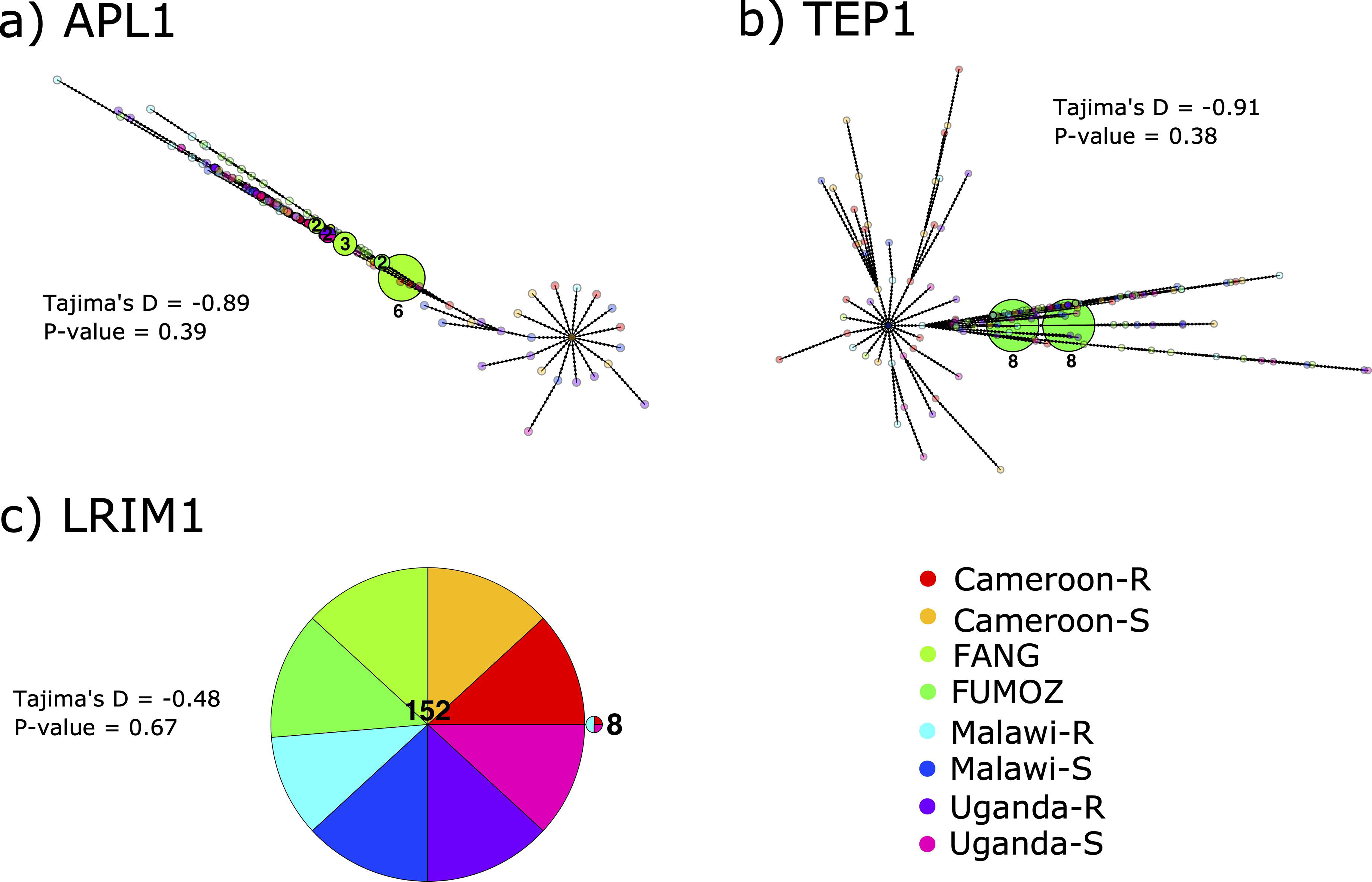
Haplotype networks of a) APL1, b) TEP1 and c) LRIM1 from SureSelect data. Haplotypes are coloured by origin (country or laboratory strain) and resistance or susceptibility of individual mosquitoes. Black circles on nodes indicate the mutational distance between haplotypes. Unique haplotypes were made translucent due to the degree of overlap along nodes. Tajima’s D and associated p-value are given for each gene adjacent to the haplotype network.

Pairwise F_st_ among PoolSeq populations revealed a slightly lower average global F_st_ for the APL1, TEP1, LRIM1 genes at 0.13, 0.12 and 0.15 respectively versus a chromosome-wide estimate of 0.16 (Supplementary Table 5), while the two gambicins had F_st_s of 0.17 and 0.13. Only six genes had global F_st_s greater than 0.3 of the 62 immune genes tested. Four of which were LRIM genes, one was TEP1 and one an APL1 paralogue (AFUN000279), however the APL1 paralogue only had two SNPs contributing to the F_st_ estimate whereas the other genes had 33-143 SNPs contributing (Supplementary Table 5). Between dead and alive SureSelect individuals, after filtering for biallelic variants 146 positions were included in the F_st_ analysis of the APL1 gene within populations. No positions were significantly different between dead and alive individuals for each population tested.

### Gambicin and other immune-related genes are also highly polymorphic

Two further immunity-related genes had elevated levels of diversity across PoolSeq populations and SureSelect data, a gambicin (AFUN006611) and a fibrinogen domain containing gene (AFUN019026) (Supplementary Table 6). The fibrinogen gene has six VectorBase paralogues, of which only one, AFUN014704, also had elevated diversity restricted to southern African populations (Supplementary Table 6). Gambicin (AFUN006611) also had a π_N_/π_s_ ratio greater than one in nine PoolSeq populations and a SureSelect π_N_/π_s_ of 1.55. This gene has two nearby gambicin paralogues in the *An. funestus* genome (AFUN006610 and AFUN006612). AFUN006610 also had an average PoolSeq π_N_/π_s_ greater than one at 1.37 while AFUN006612 did not, at a π_N_/π_s_ ratio of 0.09 (Supplementary Table 6) , AFUN006610 was not included in the SureSelect baits hence no estimate was made from these data. Haplotype networks differed in topology between the two gambicin genes (Supplementary Figure 2). AFUN006610 was dominated by one haplotype of 142/160 sequences, whereas for AFUN006611 the largest cluster was of 59 sequences with smaller clusters of up to 12 sequences. Unlike APL1 and TEP1 mutational distances between clusters were low, with at most two mutations separating clusters in AFUN006611 and only one in AFUN006610. Neither gene showed separation between origin and resistance status of included mosquitoes. Average coverages of gambicin genes were 39, 40, and 39-fold for AFUN006610, AFUN006611 and AFUN006612 respectively, which did not indicate a possible duplication event underlying elevated diversity. Other AMP genes lacked the consistency in high π_N_/π_s_ ratios of gambicins, with only a cecropin (AFUN000369) having elevated levels in the three Mozambique pools from 2002, 2016 and FUMOZ resistance strain at 1.09, 0.98 and 0.98 respectively.

### APL1 expression is greater than its paralogues and a subset of AMPs are very highly expressed

Of the five APL1 paralogues, APL1 is expressed more highly across Africa than the other genes at an average expression level of 102.43 (TPM, Supplementary Table 7). The next closest APL1 paralogue in expression is AFUN000597 at a mean TPM expression of 17.45. It is important to note however, that there may have been some artefactual expression between APL1 and paralogues in hard to discern directions as underlying gene conversion will have affected RNASeq read mapping. Indeed cross-mapping reads were used to support the inference of gene conversion as part of this study. One TEP gene (AFUN02066) syntenic with TEP15 and TEP2 in *An. gambiae*, had double the average of APL1 at 201.30 TPM. Two LRIM genes were also of higher average expression than APL1 (AFUN003917 and AFUN005964), while the TEP1 gene AFUN018758 was slightly lower at 73.54 TPM (Supplementary Table 7, per-replicate and average normalised expressions of APL/TEP and LRIM genes). For AMP genes, three cecropins (AFUN011465, AFUN015822 and AFUN015823) were expressed at by far the highest TPM of all immune genes tested (means 1448, 2691, 2031 TPM, Supplementary Table 7) followed by a defensin (AFUN006915) at 795 TPM. The two gambicin genes and their paralogue were expressed at a much lower TPMs of 14 (AFUN006610), 1 (AFUN006611) and 1 (AFUN006612) TPM respectively.

RNASeq results for APL1 revealed three contrasts of FANG, Ghana and Uganda all versus Cameroon were significant, with expression being higher in Cameroon in each case (Supplementary Table 8). For TEP1, six contrasts were significant in which Ghana and Uganda both have lower expression than Cameroon, FANG and FUMOZ. LRIM1 was significantly lower expressed in Ghana versus both Cameroon and FANG. For all three genes log_2_-fold changes were not large (< 2). For the two gambicins and paralogue only AFUN006610 was significant across ten of fifteen contrasts, and expression was highest in susceptible FANG and low in resistant FUMOZ (Supplementary Tables 7 and 8). Of the cecropin and defensin genes, AFUN011465 was the most interesting. It was significant for six contrasts including all five contrasts involving FANG where it is expressed an average of 2,340 TPM across FANG replicates versus 1,244 for all other replicates combined (Supplementary Tables 7 and 8).

### Discordant read mappings consistent with gene conversion between APL1 paralogues

We isolated discordant read-pair mappings in PoolSeq and SureSelect datasets for APL1 and its paralogues from the combined Africa-wide datasets for each. This revealed that discordant mappings of paired reads occur extensively between paralogues (Figure 3, Supplementary Figure 3). For all five paralogues discordant mappings were identified across the two exons of each gene. These mappings were biased to exon 2 for each paralog in PoolSeq and SureSelect data, which forms the majority of coding sequence across the APL paralogues. In AFUN000288 no discordant mappings were observed across exon 1 (Supplementary Figure 3). The only notable gene which experienced cross-mapping of reads to APL1 paralogues was gene AFUN020339. However, this was an artifact of paralogue AFUN000279 being encoded in an intron of this gene. APL1/AFUN018743 shared most discordant read-mappings with AFUN015851, the paralogue which also high average PoolSeq coverage.

**Figure 3.**
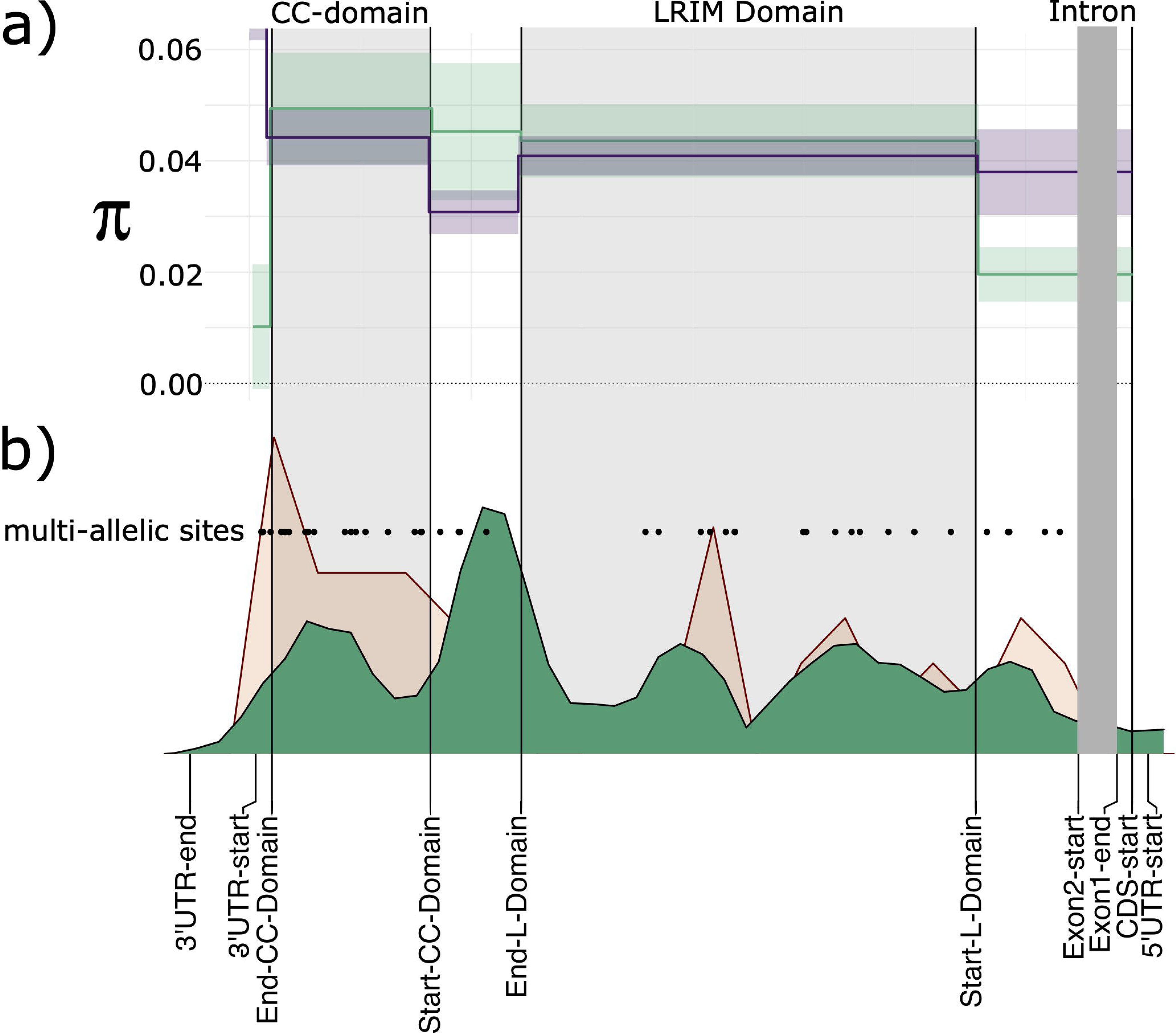
Diversity of APL1 domains and discordant read pair mappings. a) Nucleotide diversity (π) for non-synonymous sites in purple and synonymous sites in green; line is mean across twelve PoolSeq populations and shaded area is standard deviation. b) Discordant read mapping density across the APL1 gene-body is given in green in lower figure, density of multi-allelic positions is shown in salmon pink and positions of such sites are plotted as black dots labelled “multi-allelic sites”. CC-Domain is the region encoding two coiled-coil motifs and LRIM domain, both are shaded in light grey. The single intron is shaded in dark grey. Domains, exon and untranslated region positions are given on the X-axis.

By converting variant positions between APL1 paralogues to those of the multiple sequence alignment and verifying that the same reference/ancestral and alternative allele occurred at overlapping positions we identified shared polymorphisms. Shared polymorphisms remained after discordant read mappings were removed from variant predictions (Venn diagram, Figure 4). The greatest overlaps were between APL1, AFUN018581 and AFUN000279 at 25 and 20 variant positions respectively. Both GARD models predicted 20 breakpoints across the APL1 paralogue alignment, and each was significant for breakpoints versus rate heterogeneity across the alignment. The longest unbroken segment of the alignment was between positions 257-1320 of the 3075bp long alignment. This corresponded to the coding region present only in AFUN000288 but was equivalent to a length of 72 bp only in the other paralogs, a length that was consistent with other breakpoint lengths predicted by GARD (Supplementary File 1).

**Figure 4.**
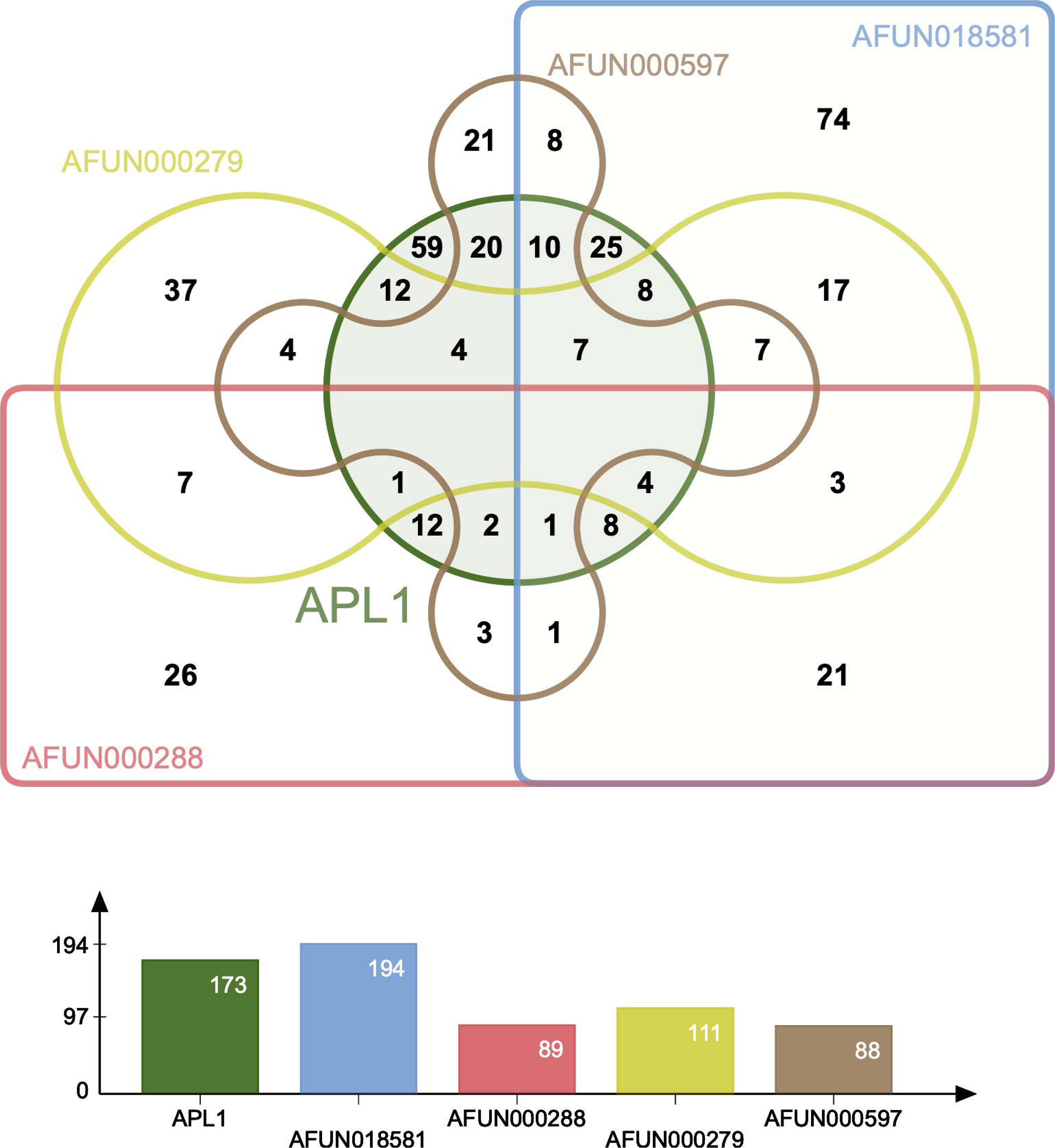
Venn diagram and bar chart of shared polymorphisms for APL1 paralogues. Polymorphisms that occur in the same alignment position of the same reference and alternate alleles between paralogues after removal of discordant read mapping. Each gene is labelled with a different colour, total number of shared positions per gene is given in the bar chart below.

Two coiled coil domains were predicted by DeepCoil in the 3’ region of all 5 paralogues (Supplementary Figure 4). In APL1, non-synonymous mutations occurred more frequently in these coiled-coil domains in PoolSeq data: 56 such sites were identified in the 313 bp covering these two domains, versus 188 across the 1,593 bp of remaining coding sequence. Furthermore, the coiled-coil domains contained 10 multi-allelic nonsynonymous SNPs, a rate twice as high (0.032 per non-synonymous site vs 0.016) as for the rest of the gene which has 25 such multi-allelic nonsynonymous variants. This coiled-coil region was not the region with the most discordant mappings in APL1 (Figure 3).

## Discussion

The *An. funestus* APL1 gene and its paralogues encode a 5’ signal peptide, LRIM domain and 3’ end coiled-coil region. This is in line with *An. stephensi* APL1 to which *An. funestus* is more closely related than *An. gambiae s. l.* (Mitri, Bischoff, Eiglmeier, et al. 2020; Neafsey et al. 2015). None of the *An. funestus* APL1 genes contain the variable ‘PANGGL’ motif of present on the 5’ end of *An. gambiae* APL1C and APL1A (Rottschaefer et al. 2011), nor does the APL1 of the more closely-related *An. stephensi* (Mitri, Bischoff, Eiglmeier, et al. 2020). None of the genes investigated here, including AMPs, had patterns of diversity restricted to their regions or resistance status (Figure 2, Supplementary Figure 2), although FANG and FUMOZ formed clusters in APL1 and TEP1 haplotype networks respectively whereas almost zero field samples had concordant haplotypes.

### Gene conversion explains elevated APL1 diversity in *An. funestus*

The three genes that form the pathogen eliminating complement system in *Anopheles* have a distinct pattern of diversity to their *An. gambiae* counterparts. Elevated APL1 is similar to that of *An. gambiae* but no population exhibits the hallmarks of a selective sweep like that of the APL1C *Plasmodium* resistance allele in *An. coluzzii* (Rottschaefer et al. 2011). Unlike *An. gambiae*, TEP diversities are not outliers compared to backdown levels species (Obbard et al. 2008), although the highest π_N_ values were elevated (>0.015) for four of these genes (Supplementary Table 3). LRIM genes exhibited the least elevated diversity of the three types, similar to LRIM genes in *An. gambiae* (Obbard et al. 2008; Slotman et al. 2007). Results consistent across PoolSeq and SureSelect. No genes from across each category appeared to have undergone a recent selection sweep, which would have been shown by reduced π_N_ and π_s_ at the affected locus and surroundings. We also found no link between insecticide resistance and allele frequencies for the principal APL1 locus (AFUN0018743). Thus, in these data a link between the insect complement system and vector competence cannot be made. By contrast increased GST expression in *An. funestus* does lead to a higher *Plasmodium* load through metabolism of parasite killing ROS and/or a fitness trade-off that decreases the robustness of immune responses (Tchouakui et al. 2019).

This observed diversity is not an artifact of gene duplication of the APL1 gene or reads mapping between paralogues due to high sequence similarity. When read-pairs that mapped between APL1 and its paralogues were removed, the genetic diversity of APL1 (π_N_ and π_s_ ) remained high in both pooled and target-enriched datasets. Furthermore, shared polymorphisms between paralogues were identified after the removal of discordant read-pairs filtered from the datasets. This is consistent with gene conversion between paralogues of APL1 that maintains high diversity at the principal locus. The paralogue alignment-based GARD approach detected many recombination events among the five APL1 paralogues which is concordant with gene conversion (Kosakovsky Pond et al. 2006). *An. stephensi* APL1 diversity sampled from Iran did not exhibit the elevated polymorphism of *An. funestus* found in all populations sampled here (Mitri, Bischoff, Eiglmeier, et al. 2020). In *An. gambiae* elevated APL1 diversity was not associated with increased interspecific divergence through relaxed constraint or higher than average mutation rate (Rottschaefer et al. 2011). In fact, lower inter-specific divergence was observed, consistent with adaptive maintenance of variation. We hypothesise that *An. funestus* will also follow this pattern when population genetic data for closely-related species is available.

The observed gene conversion-mediated increased diversity in *An. funestus* APL1 mirrors that found for the TEP1 gene in *An. gambiae* (Obbard et al. 2008). It is likely true for APL1 paralogues in *An. gambiae* as polymorphisms and the PANGGL motif are shared between loci (Rottschaefer et al. 2011). *An. gambiae s. l.* TEP1 was shown to exchange diversity with TEPs 5 and 6 through gene conversion (Obbard et al. 2008). However, high diversity in TEP1 was principally due to the presence of two divergent allele classes, one of which provides resistance to *Plasmodium* infection. When analysed separately the two allele classes exhibited diversity levels close to other *An. gambiae* loci (Obbard et al. 2008). This is similar to the *An. funestus* orthologue of TEP1, and other TEP genes which do not show elevated polymorphisms across Africa (Supplementary Table 3) suggesting no such divergence of haplotypes in this species. Although the granularity of the TEP1 haplotype network (Figure 2b) suggests processes may be occurring in this gene that are too subtle to detect by the population genetic metrics applied here.

Gene conversion can lead to increased diversification at a locus under a model in which multiple donor genes contribute diversity either reciprocally or to a recipient single locus (Kurosawa & Ohta 2011). This is not restricted to complete genes as pseudogenes of the spirochaete *Borrelin hermsii* and African Trypanosomes can also donate diversity to surface protein genes in order to evade host immune responses (Restrepo & Barbour 1994; Morrison et al. 2009). Here, we found that the APL1 *An. funestus* locus exhibits high diversity, as do its four paralogues to a lesser extent. This is a different outcome to the more frequently observed gene conversion between a single donor and recipient locus which leads to sequence homogenisation, which has occurred between the tandemly-duplicated insecticide-resistance conferring *Cyp6P9a* and *Cyp6P9b* loci in *An. funestus* from Benin (Weedall et al. 2020). Kurosawa & Ohta (2011) propose that a highly expressed gene could become the principal recipient locus among a set of paralogues experiencing reciprocal gene conversion (Kurosawa & Ohta 2011). We observed such a gene expression relationship between APL1 which is both the most highly expressed and genetically diverse gene among the five paralogues. This relationship is hypothesised to result from regulation of gene expression by local DNA accessibility, which is controlled by epigenetic modifications on surrounding chromatin. A gene more highly expressed than others in its gene conversion network will have more accessible DNA, which enhances the probability of homologous recombination occurring at this locus (Kurosawa & Ohta 2011).

The elevated diversity of APL1 has been maintained in *Anopheles gambiae s. s.* post-duplication of APL1 into three copies (Rottschaefer et al. 2011). This implies that elevated diversity is an important component of APL1 function, as a simple model of sub-functionalisation of gene-copies post-duplication, through differential pathogen targeting for example, would predict a reduction in allelic diversity at each locus.

### Elevated π_N_ in the *An. funestus* anti-microbial protein gambicin

The high gambicin gene π_N_/π_s_ ratios indicate positive selection, alternatively balancing selection could explain both π_N_/π_s_ greater than one and outright high levels of π_N_ shown in Figure 1 (Unckless & Lazzaro 2016). Alternatively, this locus could be undergoing gene conversion as is likely occurring at APL1. One test of for balancing selection would involve testing for trans-species polymorphisms at these two loci in other *funestus* group species. Such trans-species polymorphisms are commonly found in AMPs among *Drosophila* species (Unckless & Lazzaro 2016). We also note that this is a relatively short gene with ∼61 synonymous sites which may make π_N_/π_s_ ratio more susceptible to stochasticity than longer genes. Interestingly, gambicin is the only AMP among Dipterans that has been previously associated with positive selection (Lehmann et al. 2009; Arcà et al. 2014; Viljakainen 2015). Among the wider insects evidence for positive selection on AMPs is patchy, despite the natural expectation that AMPs will evolve rapidly under selection from microbes (Viljakainen 2015). This has been explained by the diversity of AMPs that insects encode, creating a dynamic environment which prevents selection on any one component (Lazzaro 2008). Rather selection is hypothesised to act on speed and efficiency of transcription and translation of AMPs post-infection (Sackton et al. 2007; Viljakainen 2015).

Current models of insect AMP, specifically cecropin, activity assume low-levels of expression in midguts, fat bodies or reproductive tracts until an immune challenge ramps up transcription (Brady et al. 2019). Our observation of three cecropins and a defensin expressed at much (>700-fold plus) higher levels than other AMPs indicate this is an incomplete model. Especially so, as expression of three of these four genes was consistent across African populations and laboratory strains which suggests constitutive high-expression of these genes in the face of diverse environmental pathogen challenges. The fourth gene, AFUN011465, a cecropin, is particularly interesting due to both its high expression and greater expression in susceptible FANG strain. We hypothesise that FANG strain mosquitoes are able to invest more resources into expression of this immune gene as they do not incur the energetic costs of metabolic resistance mechanisms versus other strains. Fitness costs of metabolic resistance on *An. funestus* are well-established with respect to life-history traits (Tchouakui et al. 2020, 2021). Indeed, the opposite has been observed in the moth *Trichoplusia ni*, in which investment in immunity has its own negative effect on developmental fitness (Freitak et al. 2007). Conversely AMP genes may be ‘cheap’ to express constitutively and therefore have little to no effect on overall fitness, as found for diptericin in *Drosophila melanogaster* (Fellous & Lazzaro 2011). Under this scenario, we would not expect a trade-off to explain higher AFUN011465 expression in susceptible FANG, although the added burden of an additional factor in resistance-conferring gene expression may make AMP expression more costly than otherwise. Linking fitness traits measured phenotypically to immune homeostasis through gene expression would further our understanding of the mechanisms underlying resistance trade-offs. This hypothesis can be tested by pathogen survival analyses on different strains of *An. funestus* versus FANG in controlled conditions without insecticide exposure. Such an experiment would require some knowledge of the pathogen range of AFUN011465 with respect to Plasmodium, bacteria and other threats.

## Conclusions

*An. funestus* has high diversity at the key immune complement factor APL1 which is most likely due to non-homologous gene conversion between five paralogues of this gene. This in line with APL1 genes in *An. gambiae s. l.* in sub-Saharan Africa, but not the more closely related *An. stephensi*. We hypothesise that open chromatin at the most expressed copy (APL1/AFUN018743) may explain preferential replenishing of this locus with diversity from other paralogues. The TEP1 gene that APL1 interacts with does not exhibit high diversity or selective sweep signals seen in *An. gambiae s. l.*, nor do numerous paralogues of this gene. The final gene of the pathogen-eradicating complex, LRIM1, also lacks a pattern of diversity different from background levels similar to other Anopheles species investigated to date. We also show that gambicin anti-microbial peptide genes have elevated nonsynonymous diversity. Gambicins are the only AMP genes that have been previously associated with positive selection in insects and gene conversion or balancing selection could explain these observations. Cecropin and defensin AMPs were constitutively highly expressed across *An. funestus* somewhat at odds with expectation, and therefore warranting further investigation.

## Data Availability

Read data for PoolSeq, SureSelect and RNASeq analyses is available in the European Nucleotide Archive under accessions PRJEB13485, PRJEB24384, PRJEB35040, PRJEB24351, PRJEB24520, PRJEB47287, PRJEB48958 and PRJEB24506.

## Supplementary Figures

**Supplementary Figure 1.** Comparison of APL1 coverage with that of a known duplicated gene. In this case read coverages showing a duplication spanning the CYP6AA1 and part of the CYP6AA2 loci in Benin first identified in (Weedall et al. 2020) and the APL1 locus also from Benin. Plots were generated in IGV (v2.8.13).

**Supplementary Figure 2.** Haplotype networks of a) AFUN006610 and b) AFUN006611 gambicin genes from SureSelect data. Haplotypes are coloured by origin (country or laboratory strain) and resistance or susceptibility of individual mosquitoes. Black circles on nodes indicate the mutational distance between haplotypes. Unique haplotypes were made translucent due to the degree of overlap along nodes. Tajima’s D and associated p-value are given for each gene adjacent to the haplotype network.

**Supplementary Figure 3. Discordant read mappings across APL1 paralogues.** Density of mappings in green, domains shaded in light grey. CC = coiled-coil domain, LR = LRIM domain.

**Supplementary Figure 4.** Coiled coil domain predictions for APL1 and paralogues.

## Supplementary Tables

**Supplementary Table 1.** Read alignments for PoolSeq and SureSelect samples generated from bam alignment files using command ’sambamba flagstat’.

**Supplementary Table 2.** Chromosomal locations of APL1, LRIM and TEP genes identified in the *An. funestus* version F3 genome adapted from VectorBase version 51 gff annotation file. APL1 and paralogues are bolded. Chromosome = chromosome of gene, Source = data source, Type = type of feature, all are protein coding genes, Start = start of gene on chromosome, End = end of gene on chromosome, Strand = gene is present on forward (+) or reverse (-) strand, Gene = gene name in VectorBase format, VectorBase description = gene annotation as assigned by VectorBase.

**Supplementary Table 3.** Gene rankings and population genetic metrics for APL/TEP/LRIM immune genes of each class investigated from PoolSeq data.

**Supplementary Table 4.** Per country π_N_ and π_S_ estimates from PoolSeq data for genome-wide averages and APL1 and paralogues.

**Supplementary Table 5.** Global F_st_ estimates for immune genes estimated from PoolSeq populations. Gene = VectorBase identifier, Function = class of gene, F_st_ = global F_st_ estimate and No of SNPs = number of SNPs contributing to F_st_ estimate.

**Supplementary Table 6.** π_N_, π_S_ and numbers of non-synonymous and synonymous sites for other immunity-related genes in *An. funestus* from PoolSeq data.

**Supplementary Table 7.** Transcript per million (TPM) counts and means for APL (bolded), LRIM,TEP and AMP genes from RNASeq sampled in Cameroon (CMR), Ghana (GHA), Malawi (MAL), Uganda (UGA), and FANG/FUMOZ laboratory strains. Mean = average of all replicates across experiment. The insecticide treatment and origin of each sample is encoded in replicate names. UNX = unexposed, PER = permethrin, DDT = Dichlorodiphenyltrichloroethane (DDT). Thus, a header containing “MAL.UNX” = a replicate sample in Malawi that was unexposed to insecticide treatment.

**Supplementary Table 8.** DESeq2 performed in iDEP differential gene expression results between countries and laboratory strains for APL (bolded), LRIM,TEP and AMP genes. Log_2_-fold change and adjusted p-value are given for every pairwise contrast made, for example “FANG-Cameroon log2FoldChange” refers to the FANG versus Cameroon contrast Log_2_-fold change. Only genes with greater than 0.5 counts per million in four or more replicates passed pre-filtering in iDEP.

## Supplementary Files

**Supplementary File 1.** Log file of GARD predicted breakpoints in the codon-aligned multiple sequence alignment of the five *An. funestus* APL1 paralogues.

