## Supplementary figures and images for "Gene conversion explains elevated diversity in the immunity modulating APL1 gene of the malaria vector *Anopheles funestus*"

### Figure S1

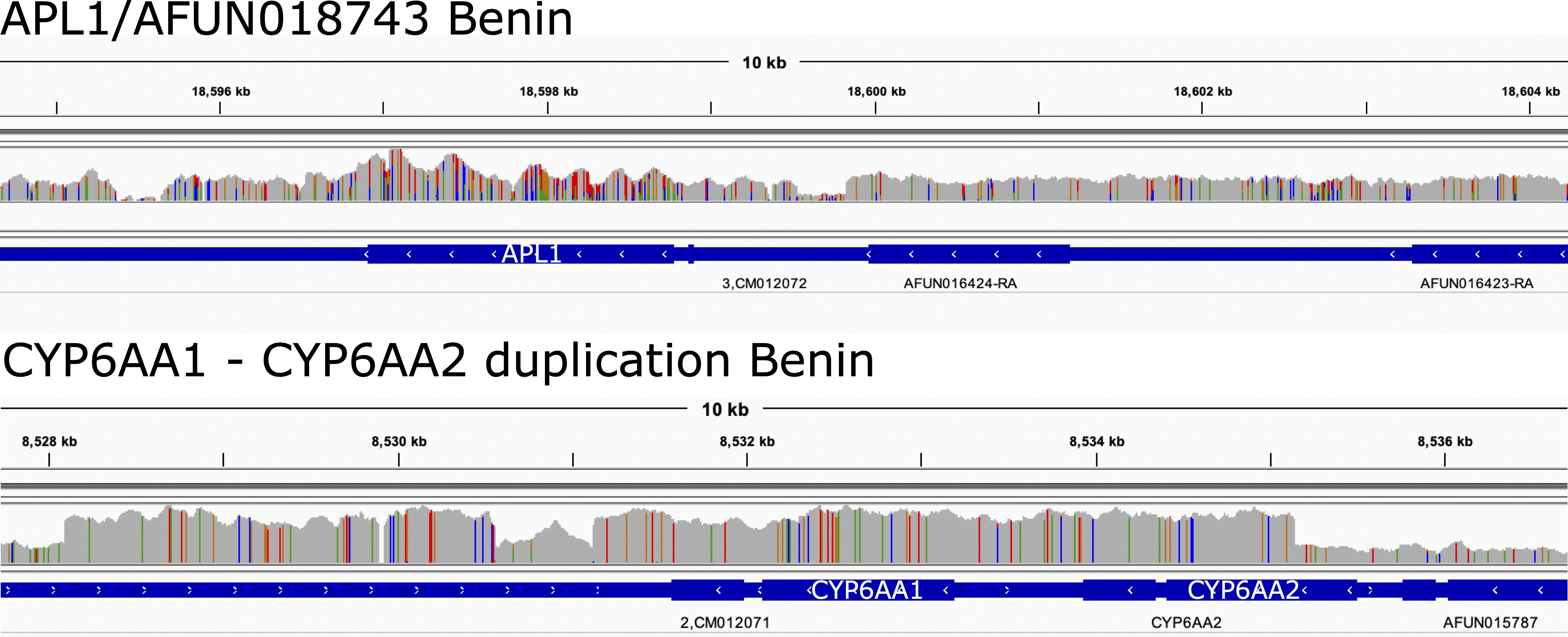

### Figure S2

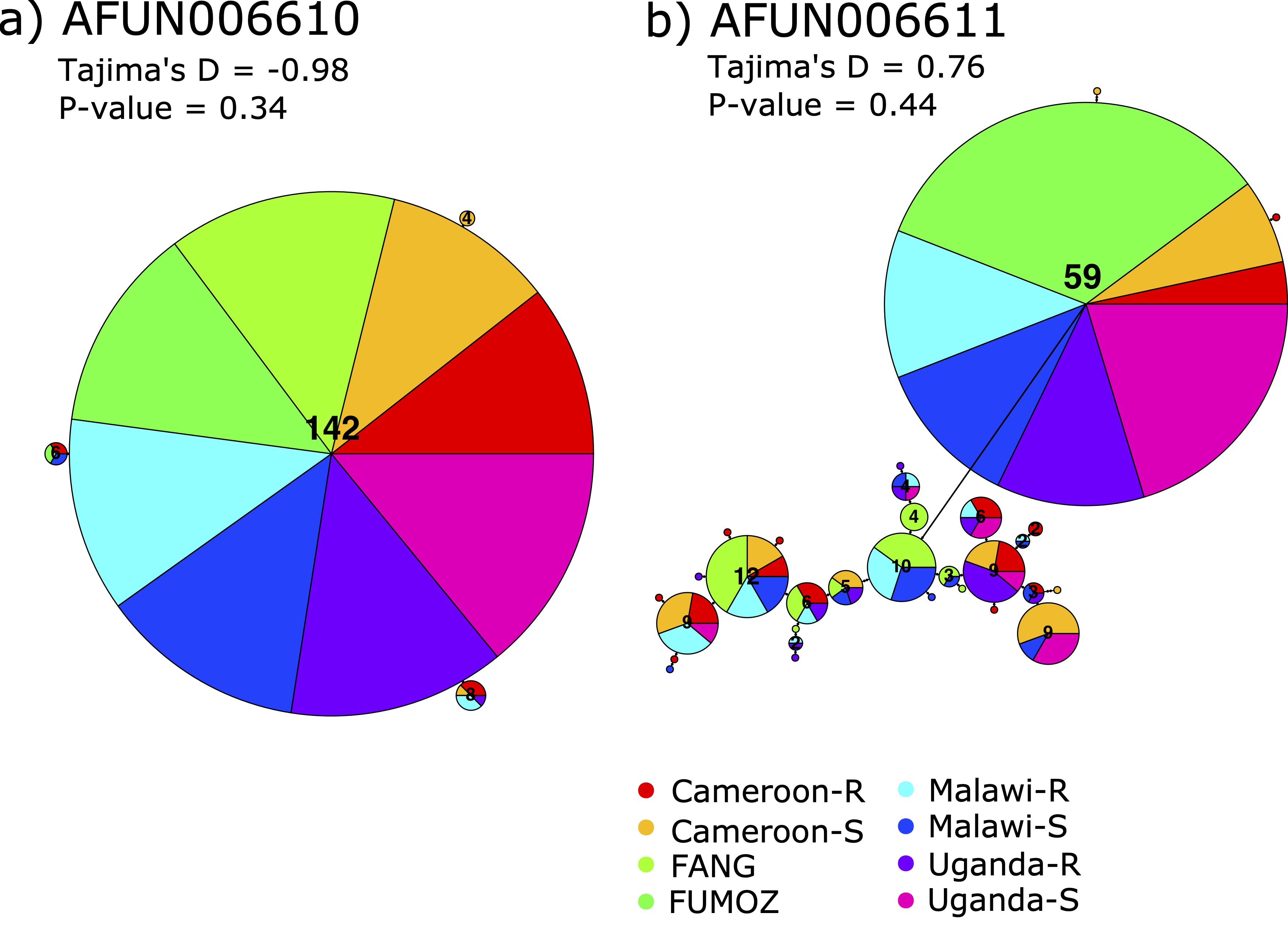

### Figure S3

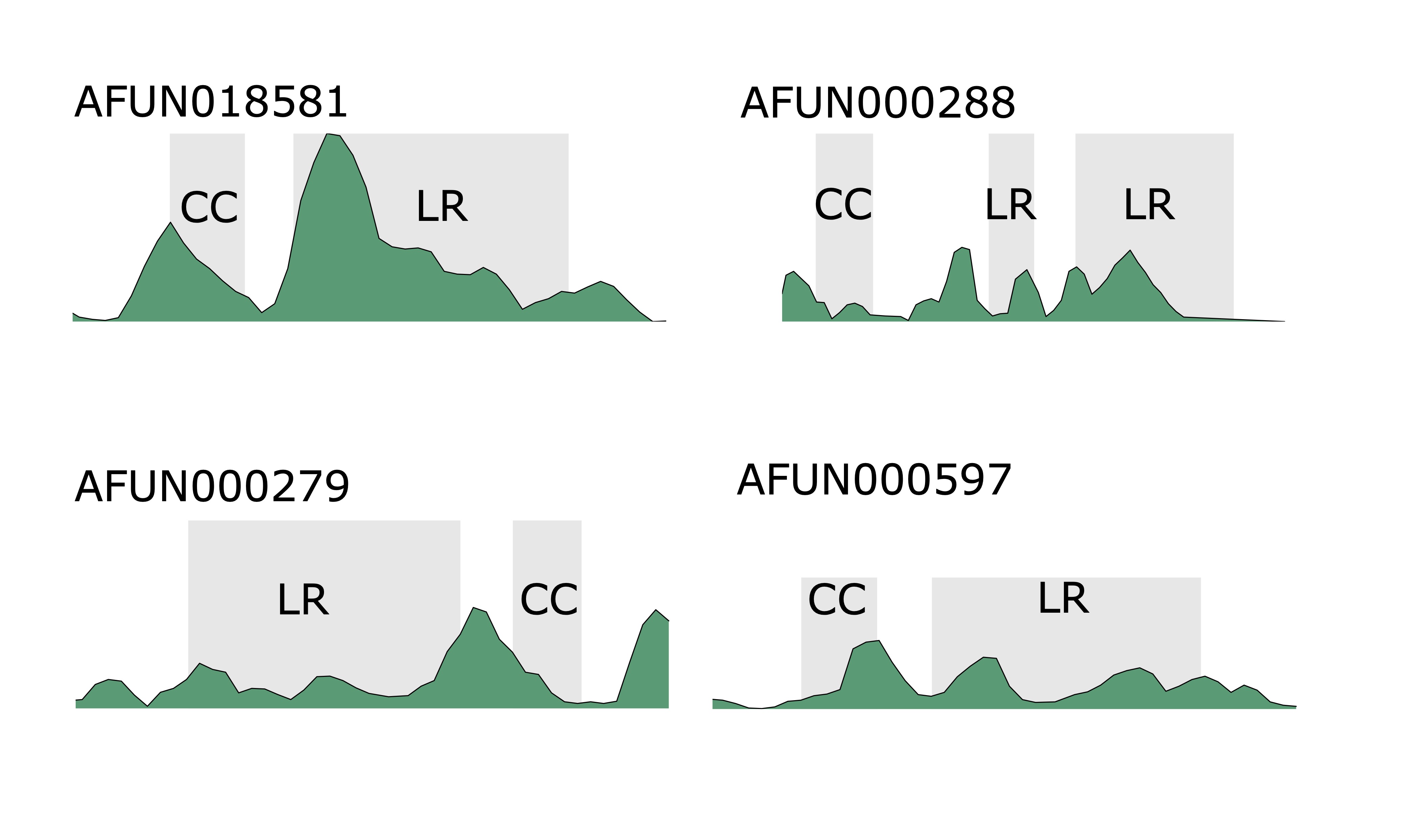

### Figure S4

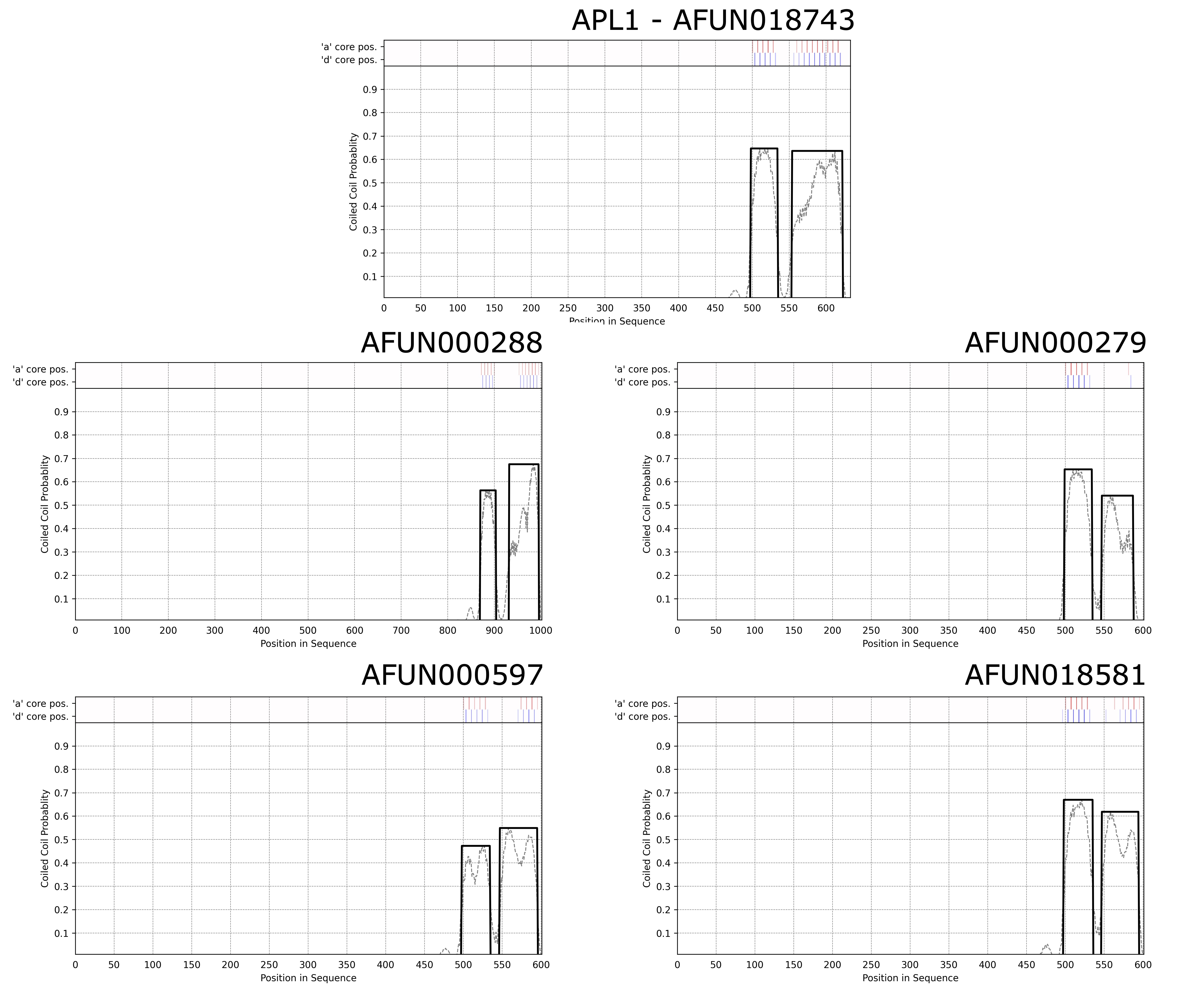
