## Supplementary material for "Gene conversion explains elevated diversity in the immunity modulating APL1 gene of the malaria vector *Anopheles funestus*": File S1

Analysis Description
--------------------
GARD : Genetic Algorithms for Recombination Detection. Implements a
heuristic approach to screening alignments of sequences for
recombination, by using the CHC genetic algorithm to search for
phylogenetic incongruence among different partitions of the data. The
number of partitions is determined using a step-up procedure, while the
placement of breakpoints is searched for with the GA. The best fitting
model (based on c-AIC) is returned; and additional post-hoc tests run to
distinguish topological incongruence from rate-variation. v0.2 adds and
spooling results to JSON after each breakpoint search conclusion

- __Requirements__: A sequence alignment.

- __Citation__: **Automated Phylogenetic Detection of Recombination Using a Genetic
Algorithm**, _Mol Biol Evol 23(10), 1891–1901

- __Written by__: Sergei L Kosakovsky Pond

- __Contact Information__:

- __Analysis Version__: 0.2

type: nucleotide
rv: None
>Loaded a nucleotide multiple sequence alignment with **5** sequences, **3075** sites (452 of which are variable) from `/home/datamonkey/datamonkey-js-server/production/app/gard/output/6012df38fced1a7bac022c34`
>Minimum size of a partition is set to be 7 sites


### Fitting the baseline (single-partition; no breakpoints) model
* Log(L) = -6959.79, AIC-c = 13949.74 (15 estimated parameters)

### Performing an exhaustive single breakpoint analysis
Done with single breakpoint analysis.
   Best sinlge break point location: 2801
   c-AIC  = 13650.3449019181

### Performing multi breakpoint analysis using a genetic algorithm
Done with 2 breakpoint analysis.
    Best break point locations: 1320, 2801
    c-AIC = 13532.34071223492
Done with 3 breakpoint analysis.
    Best break point locations: 1320, 1641, 2801
    c-AIC = 13492.46407724686
Done with 4 breakpoint analysis.
    Best break point locations: 1320, 1696, 2038, 2801
    c-AIC = 13453.21363573154
Done with 5 breakpoint analysis.
    Best break point locations: 1320, 1696, 1802, 2018, 2799
    c-AIC = 13414.77457168637
Done with 6 breakpoint analysis.
    Best break point locations: 1320, 1696, 1802, 1935, 2509, 2801
    c-AIC = 13350.83505361856
Done with 7 breakpoint analysis.
    Best break point locations: 1320, 1573, 1696, 1802, 1935, 2509, 2801
    c-AIC = 13330.53148044682
Done with 8 breakpoint analysis.
    Best break point locations: 1320, 1573, 1696, 1802, 1935, 2509, 2705, 2801
    c-AIC = 13304.47076880414
Done with 9 breakpoint analysis.
    Best break point locations: 1320, 1573, 1696, 1802, 1935, 2509, 2527, 2705, 2801
    c-AIC = 13276.36549868433
Done with 10 breakpoint analysis.
    Best break point locations: 98, 1320, 1573, 1696, 1802, 1935, 2509, 2527, 2705, 2801
    c-AIC = 13262.36679018857
Done with 11 breakpoint analysis.
    Best break point locations: 98, 256, 1320, 1573, 1696, 1802, 1935, 2509, 2527, 2705, 2801
    c-AIC = 13256.14172567599
Done with 12 breakpoint analysis.
    Best break point locations: 98, 256, 1320, 1573, 1696, 1802, 1935, 2509, 2527, 2589, 2705, 2801
    c-AIC = 13239.46552538642
Done with 13 breakpoint analysis.
    Best break point locations: 98, 257, 1320, 1531, 1573, 1696, 1802, 1935, 2509, 2527, 2589, 2705, 2816
    c-AIC = 13231.57726215827
Done with 14 breakpoint analysis.
    Best break point locations: 98, 257, 1320, 1531, 1573, 1696, 1802, 1935, 2509, 2527, 2589, 2705, 2727, 2816
    c-AIC = 13217.05832866375
Done with 15 breakpoint analysis.
    Best break point locations: 98, 257, 1320, 1531, 1573, 1618, 1696, 1802, 1935, 2509, 2527, 2589, 2705, 2727, 2816
    c-AIC = 13215.76642702358
Done with 16 breakpoint analysis.
    Best break point locations: 98, 257, 1320, 1531, 1573, 1618, 1696, 1802, 1935, 2038, 2509, 2527, 2589, 2705, 2727, 2816
    c-AIC = 13215.67941347629
Done with 17 breakpoint analysis.
    Best break point locations: 98, 257, 1320, 1531, 1573, 1618, 1643, 1696, 1802, 1935, 2038, 2509, 2527, 2589, 2705, 2727, 2816
    c-AIC = 13214.7658129462
Done with 18 breakpoint analysis.
    Best break point locations: 98, 257, 1320, 1531, 1573, 1618, 1643, 1696, 1747, 1802, 1935, 2036, 2509, 2527, 2589, 2705, 2727, 2816
    c-AIC = 13210.99045323735
Done with 19 breakpoint analysis.
    Best break point locations: 98, 257, 1320, 1531, 1573, 1618, 1643, 1696, 1747, 1802, 1827, 1935, 2036, 2509, 2527, 2589, 2705, 2727, 2816
    c-AIC = 13197.65581515156
Done with 20 breakpoint analysis.
    Best break point locations: 98, 257, 1320, 1531, 1573, 1618, 1643, 1696, 1736, 1747, 1802, 1827, 1935, 2036, 2509, 2527, 2589, 2705, 2727, 2816
    c-AIC = 13190.88784171622
